# Regulation of midzone microtubule dynamics and abscission in human cells by CAMSAP2 and Kif2a

**DOI:** 10.64898/2026.05.01.722315

**Authors:** Carline Fermino do Rosário, Erin Walsh, Andrew D. Stephens, Patricia Wadsworth

## Abstract

The spindle midzone, an array of overlapping, antiparallel microtubules, contributes to chromosome segregation and cytokinesis. As cells exit mitosis, midzone microtubules reorganize to form the midbody, the location of cell abscission. The mechanisms governing microtubule dynamics during this transition remain incompletely understood. The microtubule depolymerase, Kif2a, has been shown to contribute to midzone microtubule length control (Uehara *et al*., 2013), but how the depolymerase is regulated is not understood. Since CAMSAPs govern minus-end microtubule dynamics, we examined their role in midzone microtubule behavior. CAMSAP2, the major CAMSAP in HeLa cells, localized to the minus-ends of midzone microtubules and cells depleted of CAMSAP2, showed similar phenotypes as cells depleted of Kif2a, including elongated and bent midzones and enlarged asters. Next, we localized Kif2a in CAMSAP2-depleted cells and vice versa. CAMSAP2 remained present and extended along elongated midzone microtubules in Kif2a-depleted cells. In contrast Kif2a localization was no longer present at microtubule minus-ends but retained at plus-ends in CAMSAP2-depleted cells. In long-term live-cell movies of CAMSAP2-depleted cells abscission at the midbody was not detected, although two daughter cells formed. Markers for abscission including ESCRT-III component CHMP2A and Spastin were mislocalized, and midzone overlap zones, marked by PRC1, were extended. Together, our results demonstrate that CAMSAP2 is essential for midzone microtubule organization and dynamics, ultimately impacting cell abscission.

## INTRODUCTION

The spindle midzone is comprised of overlapping, antiparallel microtubules that extend between the segregating chromosomes in anaphase (Vukušić and Tolić, 2021; Wadsworth, 2021). In the midzone, microtubule plus-ends are located at the cell equator while the minus-ends are proximal to the chromosomes (Euteneuer and McIntosh, 1980; Mastronarde *et al*., 1993). As the nuclear envelope reforms around the segregated chromosomes and the cell transitions to telophase, the midzone microtubules become increasingly bundled and shorten, forming the midbody, the site of abscission during the final stages of cytokinesis (Guizetti and Gerlich, 2010; Fededa and Gerlich, 2012). The regulation of microtubule organization during the transition from anaphase to telophase remains incompletely understood.

Midzone microtubules play a key role in specifying the location of cytokinetic furrowing and numerous MAPs and motor proteins, which accumulate at the overlapping microtubule plus-ends, contribute to this process (Glotzer, 2009; Green, Paluch and Oegema, 2012; Kuriyama, Mullins and Skop, 2025). In contrast, less is known about the behavior of midzone microtubule minus-ends. Previous work has shown that Kif2a, a member of the Kinesin-13 family of microtubule depolymerases, contributes to midzone microtubule organization and length regulation in an Aurora B dependent fashion (Uehara *et al*., 2013). *In vitro*, Kif2a binds both microtubule ends and preferentially induces depolymerization at minus-ends, consistent with Kif2a’s role in midzone minus-end regulation (Henkin *et al*., 2023).

Microtubule minus-ends are also regulated by Calmodulin-regulated spectrin association proteins or CAMSAPs (Akhmanova and Hoogenraad, 2015; Akhmanova and Steinmetz, 2019). For example, Patronin, the Drosophila homolog of CAMSAP, binds to and “caps” microtubule minus-ends, thereby regulating minus-end depolymerization by kinesin 13 proteins (Goodwin and Vale, 2010). In anaphase, Patronin suppresses microtubule flux contributing to Anaphase B spindle elongation (Wang *et al*., 2013). In mammalian cells, two CAMSAP homologs - CAMSAP2 and CAMSAP3 - track growing minus-ends and deposit on the newly formed lattice, creating stabilized stretches that can serve as sites of non-centrosomal microtubule outgrowth (Tanaka *et al*., 2012; Jiang *et al*., 2014). In contrast, the third homolog – CAMSAP1 - tracks growing microtubule minus-ends but does not decorate the lattice (Jiang *et al*., 2014). In neurons, CAMSAP proteins play important roles in neuronal polarity, axon specification, and dendritic branching (Baas *et al*., 2016). Similarly, in polarized epithelial cells, CAMSAPs bind and stabilize non-centrosomal microtubules and maintain overall cell polarity (Tanaka *et al*., 2012). During mitosis, CAMSAP2 is required for bridging fiber assembly despite not being detectable on the spindle. Knock out of CAMSAP2 resulted in abnormally short spindles, delayed mitotic progression, slow pole-to-pole spindle elongation, and errors in chromosome segregation (Nishizawa *et al*., 2025).

Here we report novel, telophase-specific functions for CAMSAP2. We demonstrate that CAMSAP2 localizes to midzone and midbody microtubule minus-ends and, surprisingly, depletion of CAMSAP2 results in elongated midzone microtubules. In CAMSAP2-depleted cells, Kif2a is mislocalized becoming restricted to the central overlap zone and absent along midzone microtubules. Long-term, live-cell imaging reveals that, although two daughter cells form with normal timing, CAMSAP2-depleted cells show defects in the distribution of abscission machinery and PRC1 and lack a clearly recognizable abscission of midzone microtubules.

## RESULTS AND DISCUSSION

### CAMSAP2 and Kif2a contribute to midzone microtubule organization in human cells

The reorganization of microtubules in dividing Hela cells is shown in Figure 1A. Following chromosome motion in anaphase, midzone microtubules are organized in bundles which extend between the segregating chromosomes and overlap in the equatorial region (Euteneuer and McIntosh, 1980; Mastronarde *et al*., 1993; Do Rosário *et al*., 2023). As cell division progresses, the midzone transitions from a loosely organized array with extended, fan-shaped ends (third panels, top rows Figure 1A) to a more compacted, barrel-shaped structure with blunt, well-defined ends which transitions to form the midbody (last panels; Figure 1A).

**Figure 1.**
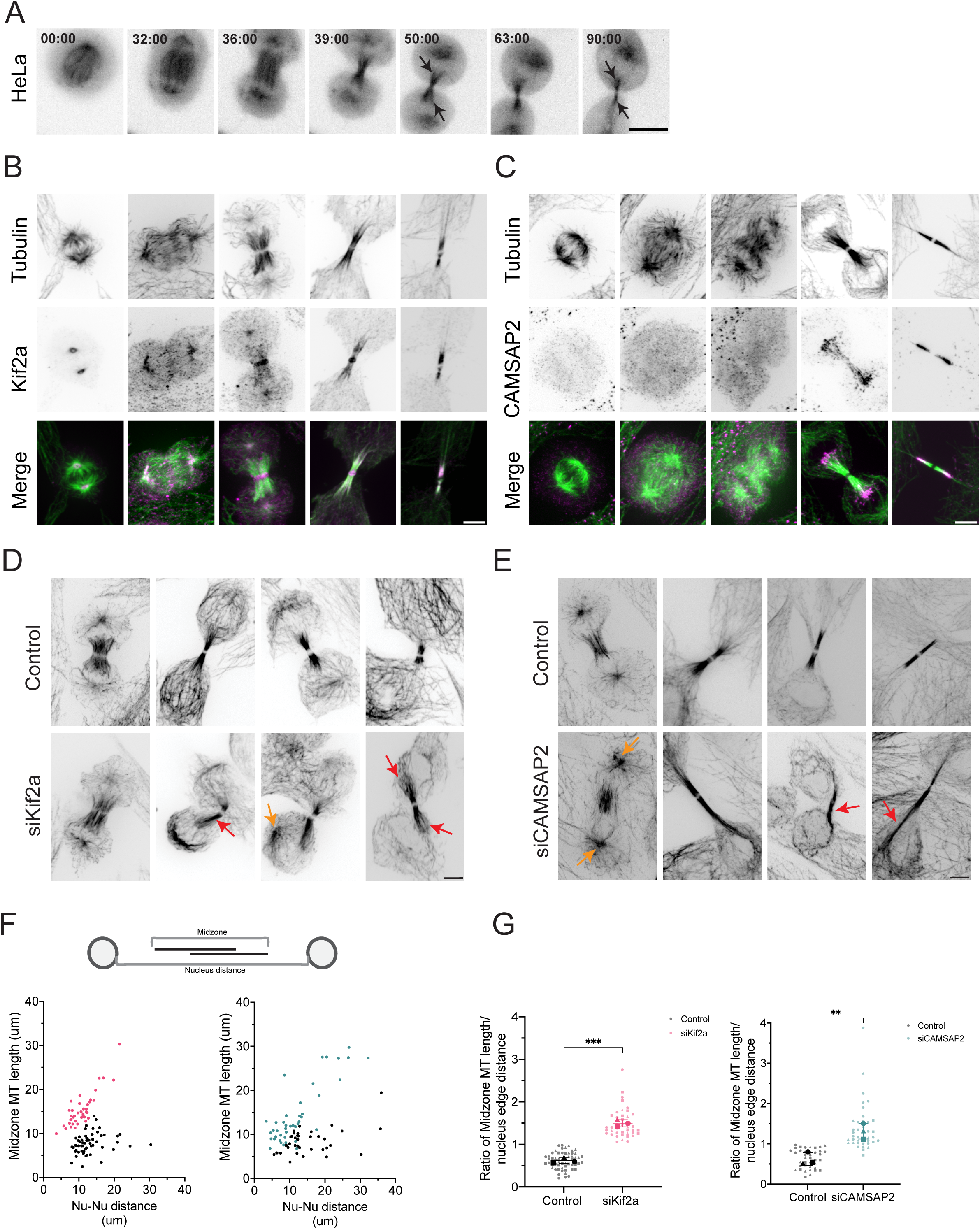
Distribution of Kif2a and CAMSAP2 in HeLa cells. (A) Microtubule reorganization during the midzone to midbody in HeLa cells expressing mCherry tubulin. Time in minutes. Immunolocalization of the kinesin-13 family members Kif2a (B) and CAMSAP2 (C) throughout mitosis. Top row shows tubulin staining; middle Kif2a or CAMSAP2; lower row shows merge (microtubules in green; Kif2a or CAMSAP2 in magenta. Organization of microtubules in control and cells depleted of Kif2a (D) or CAMSAP2 (E) using siRNA. Images depict bent, tilted, elongated midzone microtubules (shown by red arrows) and prominent astral microtubules (shown by orange arrows). (F) Diagram (top) illustrating measurements of midzone microtubules and nucleus distance. Graph showing midzone microtubule length vs nucleus distance for control and siRNA treated cells (bottom). (G) Ratio of the length of microtubule bundles/distance between nuclei was determined for control and siRNA treated cells. Each bold symbol represents the average for three trials and each symbol an individual cell; error bars represent SD. (F,G) only cells that lacked staining to Kif2a or CAMSAP2 were scored; n ≥11 cells per treatment across 3 replicates. Scale bar = 5 µm. Statistical analysis (G) Unpaired two-tailed student’s t-test; p-value legends: ns = P > 0.12; * = P < 0.033; ** = P < 0.002; *** = P < 0.001.

To understand the reorganization of midzone microtubules, we used immunofluorescence to localize both Kif2a and the minus-end binding protein CAMSAP2, potential regulators of midzone microtubule reorganization. Kif2a was detected on spindle poles from metaphase to early anaphase and in anaphase the signal was additionally detected at the central overlap region of the midzone, the location of microtubule plus-ends (Figure 1B) (Ganem and Compton, 2004). In late anaphase and telophase, Kif2a signal was detected both at the microtubule overlap and along midzone microtubules. No signal for Kif2a was detected in interphase cells (Supplemental Figure 1A).

Next, we investigated the localization of the microtubule minus-end binding protein CAMSAP2 in the midzone. Although mammalian cells contain three CAMSAP homologs (CAMSAP 1,2, and 3), CAMSAP3 is not expressed in HeLa cells and CAMSAP1 cannot be detected by immunofluorescence (Jiang *et al*., 2014). In contrast, CAMSAP2 is strongly expressed and is detected as short stretches at microtubule minus-ends in interphase cells (Jiang *et al*., 2014). Consistent with prior work, CAMSAP2 staining was not detected from metaphase to anaphase (Figure 1C) (Jiang *et al*., 2014). In telophase, however, CAMSAP2 signal was clearly detected at the minus-ends of midzone microtubules. No signal was detected at the spindle poles at any stage of mitosis. In interphase cells, CAMSAP2 decorated microtubules as characteristic ‘dashes’ along the lattice at minus-ends (Supplemental Figure 1B) (Hendershott and Vale, 2014; Jiang *et al*., 2014).

To understand the mitotic contributions of Kif2a and CAMSAP2, each protein was individually depleted (see Methods), and the organization of microtubules was examined by immunofluorescence microscopy. Depleted cells were also stained to verify knockdown of the target protein, and only cells lacking detectable staining were included in the analysis (Supplemental Figure 1C). In control cells in late anaphase, midzone microtubules terminate near the reforming nucleus and the minus-ends are loosely bundled and splayed (Figure 1D, middle top panels). Over time, midzone microtubules become highly compacted as the furrow ingresses. Additionally, the minus-ends of the microtubules gradually become less splayed, appearing blunt-ended (Figure 1D, last top panel). For Kif2a-depleted cells, no differences were observed in late anaphase compared to controls (Figure 1D, left panels). In telophase, however, defects in midzone microtubule organization were observed. One type of defect was bent midzones, which we characterized as midzones that deviate from a straight configuration, forming an angle or curve (Figure 1D, shown by a single red arrow). Midzones were also frequently tilted in the Z-axis. In addition to bent and tilted midzones, midzones were often elongated, either extending along the side or on top of the nucleus (Figure 1D, shown by two red arrows). Finally, prominent astral microtubule arrays were frequently observed in the depleted cells (Figure 1D, shown by a single orange arrow). Dividing cells could contain one or two prominent astral arrays, which were displaced to the side of the nucleus in some cells (Ganem and Compton, 2004).

Next, we looked at the organization of microtubules in CAMSAP2-depleted cells. While no difference in late anaphase was observed between control and CAMSAP2-depleted cells (Figure 1E, first bottom panel), CAMSAP2-depleted cells displayed prominent astral microtubules, bent/tilted midzones, and elongated midzone microtubules (Figure 1E, shown by red and orange arrows), similar to phenotypes observed in Kif2a-depleted cells. The elongated midzone microtubule bundles in the depleted cells appear thinner and extend around the nucleus (Figure 1E, shown by red arrows).

To quantify the midzone length in Kif2a- and CAMSAP2-depleted cells, we scored midzone microtubule organization in fixed cells, including only cells absent of Kif2a or CAMSAP2 staining, respectively. We measured the length of the midzone microtubules and the distance between the edge of each nucleus adjacent to the midzone across both control and siRNA-treated cells (Figure 1F, schematic diagram; see Methods). Both Kif2a- and CAMSAP2-depleted cells had longer midzone microtubule lengths across all nucleus-edge distances compared to control cells (Figure 1F). This was also seen when the ratio of midzone microtubule length to nucleus edge distance is calculated (Figure 1G). While elongated midzones were observed more frequently in CAMSAP2-depleted cells, fixed-cell analysis precluded determination of when these defects arose. We therefore turned to live-cell imaging.

We imaged living HeLa cells expressing mCherry tubulin that were depleted of either Kif2a or CAMSAP2. In single images of live-cells and short time-lapse sequences, we confirmed that the depleted cells showed the defects observed in fixed cells (Figure 2A). Quantification of the defects in Kif2a- and CAMSAP2-depleted cells showed that most cells contained bent or tilted midzone microtubules and enlarged astral arrays (Figure 2B). Live-cell imaging also revealed small foci of mCherry tubulin in both control and siRNA treated cells; a phenotype not observed in fixed samples (Figure 2A – C, shown by red arrows). Foci were also more frequent observed in the depleted cells, suggesting that loss of a microtubule depolymerase (Kif2a) or a minus-end binding protein (CAMSAP2) can result in additional microtubules that are not associated with the spindle or centrosomes (Figure 2B, extra foci).

**Figure 2.**
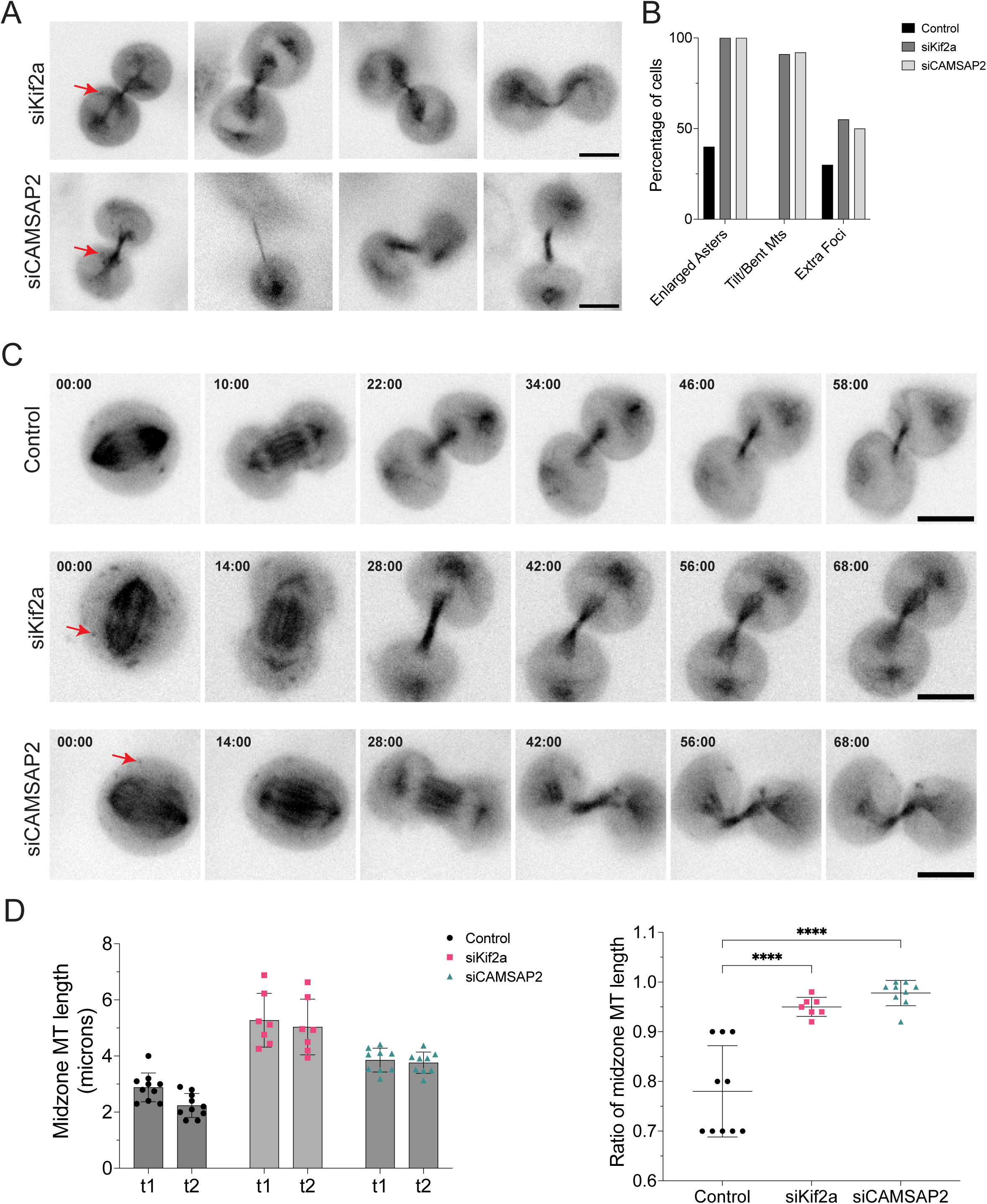
Kif2a and CAMSAP2 contribute to midzone organization and shortening in HeLa cells. (A) Individual images from timelapse of cells depleted of Kif2a (top) and CAMSAP2 (bottom). (B) Bar graph showing percentage of cells for control, siKif2a, and siCAMSAP2 for each phenotype category; n ≥ 9 cells per treatment. (C) Representative images from timelapse sequences of HeLa cells expressing mCherry tubulin; control (top) and cells depleted of Kif2a (middle) and CAMSAP2 (bottom). Time in minutes indicated in the upper left. (D) Graph of midzone microtubule length measured from two time points from movie sequences (left, see Methods). Ratio of midzone length between the two time points for control, siKif2a, and siCAMSAP2 (right); n =9 cells per treatment. Scale bar = 5 µm. Statistical analysis (D, right) Two-way ANOVA with Šídák multiple comparison test; p-value legends: ns = P > 0.12; * = P < 0.033; ** = P < 0.002; *** = P < 0.001.

Next, we acquired longer time-lapse sequences from metaphase to early telophase for control and depleted cells (Figure 2C). In control cells, midzone microtubules become compacted and shortened as telophases progressed (Figure 2C, top panels). In siRNA-treated cells, midzone microtubules remained long after ∼1 hour and, although the central region became more bundled, the ends of the midzone remained ragged (Figure 2C, middle and bottom panels). We measured the length of midzone microtubules between two time points for both control and treated cells (see Methods). The length of the midzone decreases between the two time points for control cells, although the change in length is minimal (Figure 2D, right). However, we see no change in the midzone microtubule length between the two time points for the treated cells (Figure 2D, right). The ratio of midzone microtubule length (t2/t1) is higher for siRNA treated cells compared to controls (Figure 2D, left). It is also noted that the length of the midzone for treated cells at t1 is statically longer than the midzone of control cells at t1 (data not shown). These results confirm that Kif2a contributes to the regulation of midzone microtubule length and organization (Uehara *et al*., 2013) and demonstrate for the first time, that CAMSAP2 binds to and regulates microtubule minus-ends during telophase.

Previous *in vitro* studies report that CAMSAP2 and CAMSAP3 alter microtubule minus-end dynamics by reducing minus-end growth and suppressing catastrophe and thus stabilizing the minus-ends (Hendershott and Vale, 2014b; Jiang *et al*., 2014). In epithelial cells, free microtubule minus ends predominantly pause but can also undergo periods of slow growth. Loss of CAMSAP2 abolished this behavior, promoting minus-end shrinkage (Jiang *et al*., 2014). We show that depletion of CAMSAP2 resulted in elongated midzone microtubules in telophase, which is inconsistent with the reduction in shortening and the increase in catastrophe observed in interphase cells lacking CAMSAP2 suggesting that additional factors may regulate midzone minus-ends *in vivo*.

### CAMSAP2 contributes to proper localization of Kif2a at microtubule minus-ends

Because Kif2a and CAMSAP2 are both present and partially overlap in their distribution in the midzone of telophase cells (Figure 1 B,C), we investigated how the depletion of one protein impacted subcellular localization of the other. First, we looked at CAMSAP2 localization in Kif2a-depleted cells using immunofluorescence. In interphase cells, the number of CAMSAP2 dashes increased in Kif2a-depleted cells compared to controls, consistent with prior work (Guan, Hua and Jiang, 2023) (Supplemental Figure 2A). In telophase control cells, we observed that midzone microtubules shortened with their minus-ends aligned and localized away from the nucleus. CAMSAP2 staining appeared proximal to the minus-ends of the microtubules; staining did not extend completely to the middle of the midzone (Figure 3A, left panels). As seen previously, Kif2a-depleted cells displayed bent and elongated midzone microtubules that wrapped around the nucleus. CAMSAP2 signal was detected along midzone microtubules including the extended, elongated midzone microtubules in the Kif2a-depleted cells (Figure 3A, right panels).

**Figure 3.**
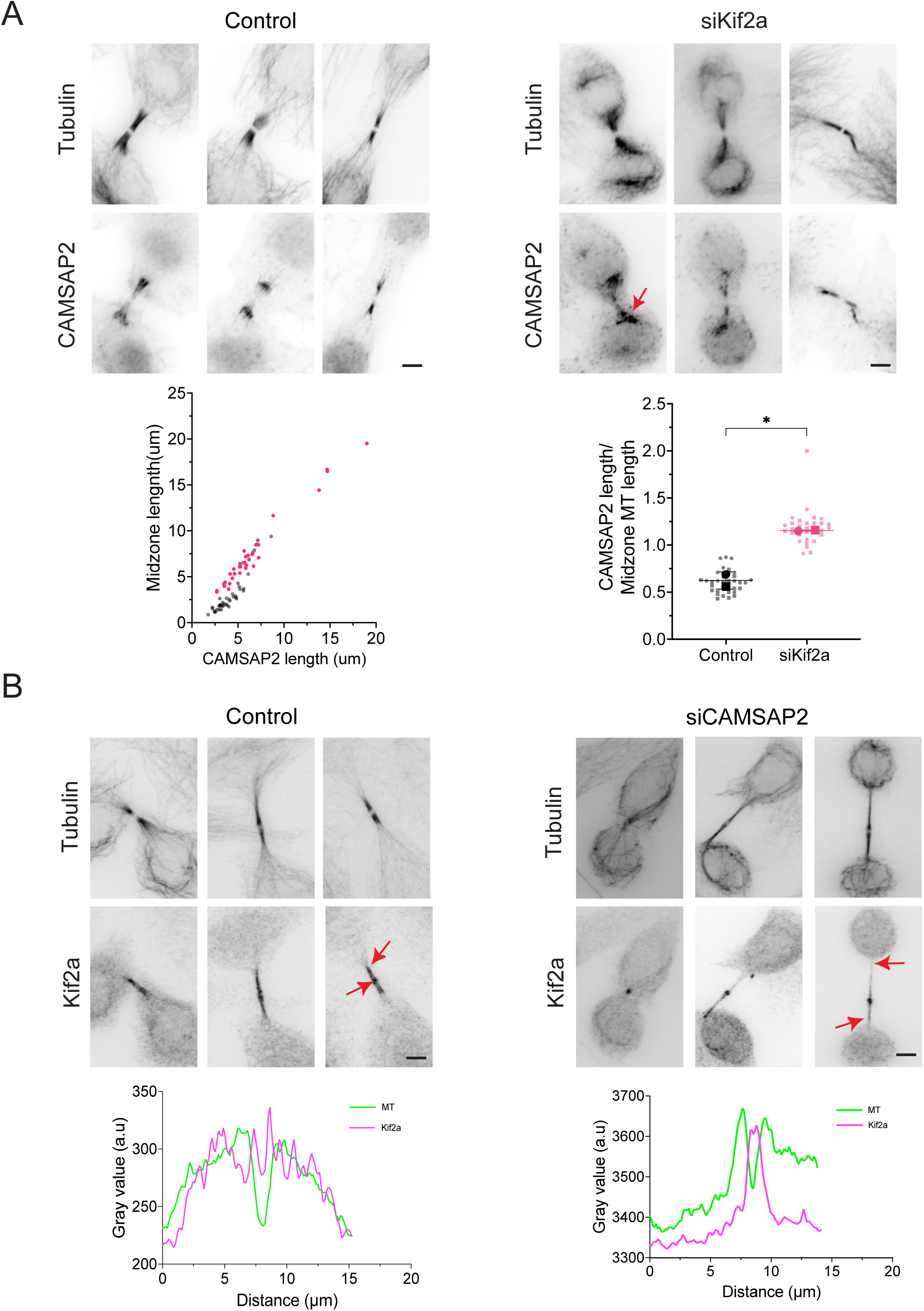
CAMSAP2 contributes to proper localization of Kif2a at microtubule minus-ends. (A) CAMSAP2 localization in control and Kif2a-depleted cells. Top row: alpha tubulin; lower row: CAMSAP2 staining. Note the more extensive CAMSAP2 distribution in Kif2a-depleted cells (shown by red arrow). Graph showing midzone microtubule length vs length of CAMSAP2 staining for control and Kif2a-depleted cells (left). Ratio of CAMSAP2 staining to microtubule length for control and Kif2a-depleted cells (right); each bold symbol represents the average for two trials and each symbol an individual cell; error bars represent SD, n ≥11 cells per treatment across 2 duplicates. (B) Kif2a localization in control and CAMSAP2-depleted cells. In control cells, Kif2a localizes along midzone microtubules and at the central microtubule overlap zone of the midbody (lower right). In depleted cells, Kif2a is not detected along microtubules (red arrows) but retains localization to the overlap. Fluorescence intensity profiles of microtubules (green) and Kif2a (magenta) signal in control and CAMSAP2-depleted cells. Scale bar = 5 µm. Statistical analysis (A, bottom right graph) Unpaired two-tailed student’s t-test; p-value legends: ns = P > 0.12; * = P < 0.033; ** = P < 0.002; *** = P < 0.001.

To determine if loss of Kif2a significantly alters CAMSAP2 cellular distribution, we measured the length of midzone microtubules and length of CAMSAP2 staining along the microtubules. Results show longer lengths of CAMSAP2 decoration (Figure 3A, pink dots, siKif2a) compared to control cells (Figure 3A, black dots, Control), tracking with the increased midzone length (Figure 3A, left plot). Calculating the ratio of CAMSAP2 length to midzone microtubule length also revealed a significant increase in Kif2a-depleted cells compared to controls.

We then investigated the effect of CAMSAP2 depletion on Kif2a distribution. In control telophase cells, Kif2a localized along midzone microtubules and at the central microtubule overlap zone of the midbody (see Figure 1 and Figure 3B, indicated by red arrows in the third control panel). To our surprise, in CAMSAP2-depleted cells, despite the presence of elongated and bent microtubules, Kif2a signal is not detected along the microtubules or toward the minus-ends but is retained on microtubules of the central overlap zone (Figure 3B, red arrows). The change in the distribution of Kif2a can be seen from the fluorescence intensity profiles for microtubules (MT, green line) and Kif2a (Kif2a, magenta line). In control cells, Kif2a intensity closely tracks with MT intensity across the midzone, showing clear co-localization (Figure 3B, left graph). In contrast, Kif2a signal in CAMSAP2-depleted cells remained at the central overlap zone but was significantly reduced along the microtubules (Figure 3B, right graph).

These results show that CAMSAP2 occupies a greater proportion of the elongated midzone microtubules in telophase in Kif2a-depleted cells. This is consistent with the observation that CAMSAPs are deposited along the minus-end proximal lattice during minus-end elongation (Jiang *et al*., 2014). However, in CAMSAP2-depleted cells, we showed that Kif2a was solely confined to the central overlap zone and was absent at the minus-ends and along microtubules. *In vitro*, Kif2a selectively depolymerizes microtubule minus-ends and in cells an Aurora B phosphorylation gradient restricts Kif2a activity to minus-ends of midzone (Uehara *et al*., 2013; Henkin *et al*., 2023), suggesting that CAMSAP2 plays a key role in regulating the activity and location of Kif2a in telophase. The coiled-coil domains of CAMSAPs have been shown to interact with katanin and kinesin-14 KIF3C, to regulate interphase and neuronal microtubules (Jiang *et al*., 2014, 2018; Cao *et al*., 2020; Liu *et al*., 2021). It is not yet known if Kif2a interacts directly or indirectly with CAMSAP2 to regulate microtubule minus-end localization and activity. Interestingly, our results also shows that plus-end binding of Kif2a is not dependent on CAMSAP2.

### Long-term imaging of CAMSAP2-depleted cells show defects in mitotic timing and abscission

Our short-term live-cell imaging showed that midzone microtubules remain long for approximately 60 minutes post-metaphase (Figure 2A) and that Kif2a minus-end localization is lost in CAMSAP2-depleted cells (Figure 3B). However, it is still unknown whether these organizational changes influence the final steps in cell division, including abscission. Because CAMSAP2 depletion affected the minus-end localization of Kif2a, we performed long-term imaging of CAMSAP2-depleted cells in which CAMSAP2 is reduced and Kif2a diminished from midzone microtubules (see Methods). Time-lapse image sequences of control and treated cells show that cells progressed through mitosis, completing the transition from nuclear envelope breakdown (NEBD) through metaphase and anaphase. Control cells took an average of 28 minutes from NEBD to metaphase and 37 minutes to anaphase, compared to 44 and 54 minutes, respectively, in CAMSAP2-depleted cells. These results reveal a statistically significant increase in the duration from NEBD to metaphase and anaphase in CAMSAP2-depleted cells (Supplemental Figure 3A,B). A recent study also reported a delay in mitotic progression, showing that the duration from anaphase onset to completion of chromosome separation was longer in CAMSAP2 knockout cells (Nishizawa *et al*., 2025).

In control cells, midzone microtubules bundle and shorten as telophase progresses, ultimately forming a compact midbody that undergoes abscission – the final severing of the intercellular bridge that physically separates the two daughter cells (Figure 4A, top panels shown by red arrow). Interestingly in CAMSAP2-depleted cells, no clear cut of the midzone microtubules was observed. Rather, the microtubules between the two daughter cells became dimmer and subsequently two daughter cells lacking any intervening microtubule structure were observed (Figure 4A, bottom panels shown by red arrow). In control cells, nearly 100% of the cells had an observable cut event (Figure 4B). In contrast, in CAMSAP2-depleted cells no examples of a cut midzone were observed. In a small percentage of control cells, two daughter cells failed to form; this percentage was greater in the depleted cells (Figure 4B).

**Figure 4.**
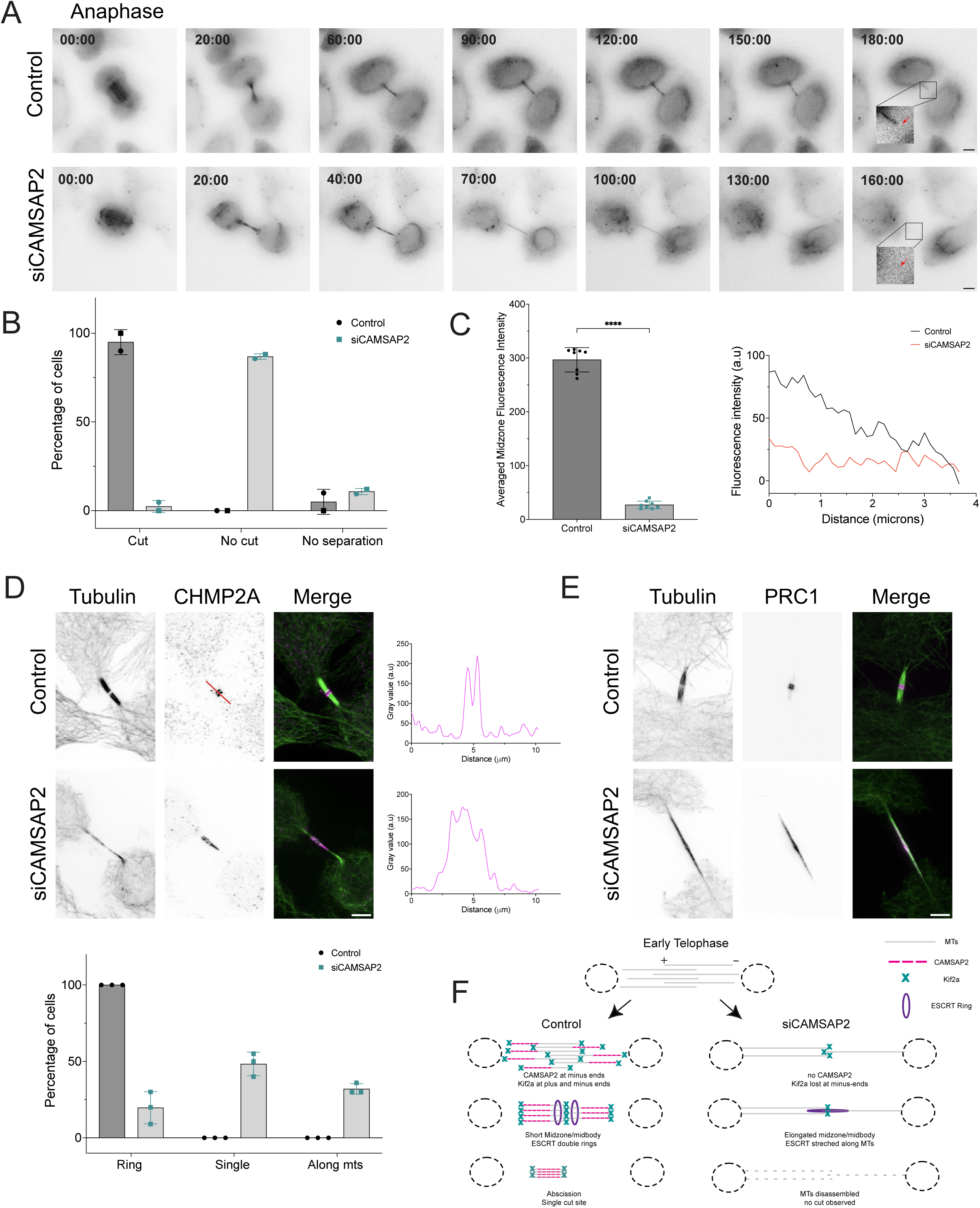
Long-term, live-cell imaging of HeLa cells expressing mCherry-tubulin. (A) Control (top row) and CAMSAP2-depleted cells (lower row) from anaphase to daughter cell separation. Anaphase is set as time zero. Time in minutes. Higher magnification views of the region enclosed by black box are shown for control and treated cell. (B) Percentage of cells displaying complete cut (successful abscission), no cut (MT fade), and no separation (failed abscission) for control and CAMSAP2-depleted cells; each symbol represents the percentage for two trials, n ≥21 cells per treatment across 2 duplicates. (C) Right: Graph of average midzone intensity for control and CAMSAP2-depleted cells; n= 8 cells per treatment across 2 duplicates. Left: Representative line profile of abscission event for control (black line) and CAMSAP2-depleted cells (red line). (D) Localization of CHMP2A in control and CAMSAP2-depleted cells (left). Fluorescence intensity profile of CHMP2A in control and CAMSAP2-depleted cells (right). Percentage of cells displaying CHMP2A localization pattern for control and CAMSAP2-depleted cells (bottom); each symbol represents the percentage for three trials; n ≥ 5 cells per treatment across 3 replicates. (E) Localization of PRC1 in control and CAMSAP2-depleted cells. (F) Model. Scale bar = 5 µm. Statistical analysis (C) Unpaired two-tailed student’s t-test; p-value legends: ns = P > 0.12; * = P < 0.033; ** = P < 0.002; *** = P < 0.001.

To understand the failure in abscission, we measured the average midzone fluorescence intensity in CAMSAP2-depleted cells and showed that it is significantly lower than control cells, indicating a decrease in microtubule density in the treated cells (Figure 4C, left graph). For control cells, fluorescence intensity profiles revealed that upon abscission one side of the midbody showed a decrease in fluorescence, but the other side remained bright (Figure 4C, black line, inset in 4A). For CAMSAP2-depleted cells, the fluorescence intensity was uniform along the midzone; no abrupt loss of fluorescence intensity was observed. These observations were consistent with the disassembly of midzone/midbody microtubules without a discrete cutting event (Figure 4C, red line). Although CAMSAP2-depleted cells lacked a discrete cutting event, the timing of daughter cell separation was not significantly different from controls (Supplemental Figure 3C).

Because no clear cut of the midzone microtubules was observed in CAMSAP2-depleted cells, we next asked if components of the abscission machinery are present in these cells. The Endosomal Sorting Complex Required for Transport, ESCRT, is required for the final abscission (Carlton and Martin-Serrano, 2007; Morita *et al*., 2007). The ESCRT-III complex directly interacts with the microtubule severing enzyme Spastin to coordinate microtubule removal (Yang *et al*., 2008; Connell *et al*., 2009; Advedissian, Frémont and Echard, 2024). Charged multivesicular body protein 2A (CHMP2A) is a core component of the ESCRT-III complex (Kamenetsky *et al*., 2025).

We first compared CHMP2A localization in control and CAMSAP2-depleted cells using immunofluorescence. In control cells, CHMP2A forms a characteristic double ring pattern at the midbody, consistent with prior work (Elia *et al*., 2011; Guizetti *et al*., 2011; Kamenetsky *et al*., 2025) (Figure 4D, top panels). In CAMSAP2-depleted cells, the localization of CHMP2A was either a single globule, a double ring or an extended pattern (Figure 4D, bottom panels). The distribution of CHMP2A can be seen from fluorescence intensity profiles of the double ring pattern (control Figure 4D, top right) and the extended pattern (Figure 4D, bottom right). These phenotypes were quantified for control and CAMSAP2-depleted cells (Figure 4D, bottom graph). All the control cells displayed a double ring pattern; in contrast, in CAMSAP2-depleted cells only 20% of cells showed double ring and the remainder showed either the extended pattern or a single globule. Since the ESCRT-III machinery recruits Spastin to the midbody, we hypothesized that Spastin localization would also be affected by CAMSAP2 depletion. In control cells, nearly all cells displayed the double ring pattern of Spastin (Connell *et al*., 2009) (Supplemental Figure 3D). In CAMSAP2-depleted cells, Spastin localization exhibited all three phenotypes, like the CHMP2A (Supplemental Figure 3D).

Since both proteins displayed defects in midzone localization, we investigated whether the midzone overlap zone is disrupted in CAMSAP2-depleted cells. PRC1, a key regulator of cytokinesis, selectively crosslinks antiparallel microtubules and becomes highly enriched at the spindle midzone overlap during anaphase and telophase (Mollinari *et al*., 2002; Kurasawa *et al*., 2004; Vernì *et al*., 2004; Zhu and Jiang, 2005). In control cells, PRC1 is concentrated at the center of the midbody (Figure 4E, top panels). However, in CAMSAP2-depleted cells, PRC1 displayed an elongated localization, indicating that the microtubule overlap zones were extended (Figure 4E, bottom panels).

These results showed that CAMSAP2-depleted cells ultimately form two daughter cells but do so without the canonical abscission event observed in control cells. In control cells, abscission was visible as a discrete cut of the intercellular bridge, whereas in CAMSAP2-depleted cells, the midzone microtubules disassembled without a detectable severing event, suggesting that the normal mechanism of microtubule removal at the intercellular bridge was disrupted. The midzone fluorescence intensity was dramatically reduced in CAMSAP2-depleted cells compared to controls, showing that fewer microtubules are present at the intercellular bridge. Strikingly, both CHMP2A and Spastin - proteins of the abscission machinery - showed altered localization in CAMSAP2-depleted cells.

We propose that CAMSAP2 regulates microtubule minus-end behavior during telophase to establish an organized midzone that supports proper Kif2a localization and the correct assembly of the abscission machinery (Figure 4F). In control cells, the midzone compacts into a short midbody with CAMSAP2 (pink dashes) decorating the minus-ends and Kif2a (teal crosses) present at both plus- and minus-ends. The ESCRT-III double rings (purple ovals) are visible at central overlap of the midbody (Figure 4F, left). In the absence of CAMSAP2, midzones are longer, Kif2a is lost from minus-ends, the abscission machinery is mislocalized, and the microtubule overlap is elongated. Although two daughter cells form with similar timing to control cells, canonical abscission is not detected. Instead, microtubule density is greatly reduced until microtubules are no longer detected (Figure 4F, right).

The mis-localization of CHMP2A, Spastin and PRC1 at the midbody in CAMSAP2-depleted cells further shows that disruption to microtubule minus-end behavior alters microtubule organization throughout the midzone/midbody. Finally, we show that cells lacking CAMSAP2 produce two daughter cells, but they do so without the canonical abscission event.

## Materials and Methods

### Cell Culture

Parental HeLa and HeLa cells expressing mCherry-α-tubulin and CRISPR EGFP-Eg5 (Mann and Wadsworth, 2018) were grown in DMEM (Sigma Aldrich, D7777) with 10% fetal bovine serum (CPS Serum, FBS-500-HI) and antibiotic-antimycotic at 37℃ with 5% CO_2_, in a humid atmosphere. For live-cell imaging experiments, the growth medium was removed and replaced with non-CO_2_ medium lacking sodium bicarbonate and containing HEPES (Sigma Aldrich, H3784). For transfections, cells were incubated in DMEM without antibiotic – antimycotic.

### Immunofluorescence

HeLa (parental or cells expressing mCherry-α-tubulin and CRISPR EGFP-Eg5 cells (Mann and Wadsworth, 2018) were plated on #1.5 22x22mm microscope cover glass for 24 - 48 hours prior to fixation. Cells were rinsed two or three times in room temperature calcium and magnesium free PBS and then fixed in -20℃ methanol for 10 minutes. Cells were rehydrated in PBS containing 0.1% Tween 20 and 0.02% Sodium Azide (PBS-TW-Az). For some antibodies, cells were rinsed in PBS and then fixed in freshly prepared 0.25% glutaraldehyde, 2% paraformaldehyde, 0.5% Triton X 100, prepared in PBS. Cells were incubated in the manufacturer’s recommended dilution of primary antibody prepared in PBS-Tw-Az; 2% BSA was added to block non-specific staining. The following primary antibodies were used: alpha-tubulin (mouse tubulin DM1a [Sigma], 1:100; rat tubulin YL ½ [Thermo Scientific], 1:100), Kif2a (Novus, 1:200; and ProteinTech, 1:200), CAMSAP2 (ProteinTech, 1:100; and Fisher Scientific, 1:500), Spastin (Santa Cruz Biotechnology. 1:100), and CHMP2A (ProteinTech, 1:100). Incubations with primary antibodies were performed in a humid chamber for one hour at 37℃ or overnight at 4℃. Coverslips were rinsed by dunking the coverslips at least 30 times in a beaker containing PBS-TW-Az. Secondary antibody incubations were also performed in a humid chamber at room temperature for 45 minutes, at the manufacturers recommended dilution, followed by rinsing as described above. The following secondary antibodies were used: Goat anti-mouse Dylight 488 (1:400), anti-mouse Cy3 (1:400), Goat anti-rabbit Cy3 (1:400), Donkey anti-rabbit 488 (1:500), Donkey anti-rat 488 (1:400), and Donkey anti-rat 555 (1:400). All secondary antibodies are from Jackson Immunoresearch Laboratories, expect Donkey anti-rat 555 (Invitrogen). Coverslips were mounted in Fluoromount G (Southern Biotech) and sealed with nail polish.

### siRNA

For delivery of siRNA, HeLa cells (parental, or cells expressing mCherry-α-tubulin and CRISPR EGFP-Eg5) were plated at a density of 250,000 cells per well in a 6-well plate on day one. The following day, the RNAi duplex-Lipofectamine RNAiMax complex was prepared for each well according to the manufacturer’s recommendations. The cells and siRNA were incubated for 24 hours at 37℃ in a 5% CO_2_ incubator. The cells were then re-seeded either on coverslips or in glass bottom dishes. All siRNAs were obtained from Sigma-Aldrich; siRNA was resuspended in the RNAi buffer to a concentration of 20uM. The sequences used for siRNA are: Kif2a 5’ATACAATGGTTCAGCTATA3’; CAMSAP 2 5’ GTACTGGATAAATAAGGTA3’; and Katanin p60 5’GGCTCGATTTTATTCTCCA3’. For control, Mission siRNA Universal Negative Control #1 (SIC001) was used.

### Imaging

For imaging fixed cells, a Nikon Ti-E microscope equipped with a 100X phase, NA 1.4 oil immersion objective, CSU-X1 Yokogawa Spinning disc confocal scan head and with a Hammamatsu ORCA-ER camera. Complete Z-stacks were collected using a 0.30 μm spacing; exposure time varied depending on the antibody. Image acquisition was controlled by Metamorph software (Molecular Devices, LLC).

HeLa cells expressing mCherry-α-tubulin were imaged using a Nikon Eclipse Ti-S microscope equipped with a 40X air objective, NA 0.7, QImaging QICAM Fast 1394 Camera, Sutter Instrument Lambda SC Smart Shutter Controller, and a temperature controller TC-500 (20/20 Technologies). Image acquisition was controlled by ImageJ software and Micromanager. Live-cells were imaged with 2-minute intervals.

For long-term imaging, HeLa cells expressing mCherry-tubulin were imaged with Nikon Elements software on a Nikon Ti2-E microscope, Orca Fusion Gen III camera, Lumencor Aura III light engine, MC CleanBench air table, equipped with a 40x air objective. Live-cell timelapse images were acquired using the Nikon Perfect Focus System. For overnight imaging, the growth medium was removed and replaced with FluoroBrite DMEM (ThermoFisher Scientific) with FBS. The cells were maintained at 37℃ at 5% CO_2_ using a Okolab heat, humidity, and CO_2_ stage top incubator (H301). mCherry-tubulin fluorescence was imaged using a 546 nm laser at 2% laser power with 30 ms exposure. To visualize the DNA, cells were incubated with Spy650-DNA (1:1000 dilution) for 30 minutes to 1 hour at 37℃ in a 5% CO_2_ incubator. The Spy650-DNA was removed before imaging. Spy650-DNA dye was imaged using a 638 nm laser at 2% with 30 ms exposure. Images were acquired at 10-minute intervals. At each interval, 5 Z-slices at 2.25 μm spacing were acquired.

### Image analysis

The length of the midzone microtubules was measured between two selected time points, chosen as follows. The timelapse was played to completion. The frame where the midbody was compacted and had stopped shortening was selected as time point 2. The movie was then played in reverse. Time point 1 was defined as the frame before shortening starts, where the midzone microtubules were already compacted but the ends still appeared fan-shaped. For the ratio, the length of the second timestamp was divided by the length of the first timestamp. This was done for both control and siRNA treated cells.

For the midzone length to nucleus distance ratio, the length of midzone microtubules was measured using a segmented line. The distance between the closest edge of the two nuclei was measured along the same trajectory used to measure the microtubules. The ratio was obtained by dividing the microtubule length by the distance between the edge of each nucleus. These measurements were obtained for control, CAMSAP2- and Kif2a-depleted cells.

The CAMSAP2 length to midzone microtubule length ratio was determined by measuring the length of the microtubules using either a straight line or segmented line on Sum Projections on only one half of the midzone. Then, the length occupied by CAMSAP2 was divided by the midzone microtubule length. These measurements were obtained for both control and Kif2a-depleted cells.

The number of CAMSAP2 dashes was determined for both control and Kif2a-depleted cells. To do this, we counted the dashes in two selected locations within an interphase cell, using a rectangular box (width = 51 px, height = 59 px) placed based on the microtubule channel. The same ROIs were then applied to the CAMSAP2 channel for dash counting. Measurements were performed on Sum Projections. For control and Kif2a-depleted cells, the average number of dashes from the two ROIs was calculated.

For the intensity plot profiles, a line was drawn along the midzone microtubules from one end to the other using the tubulin channel. The same line was then applied to the Kif2a channel. This was done for both control and CAMSAP2-depleted cells.

For the CHMP2A distribution, a line was drawn along the staining using the CHMP2A channel for both control and CAMSAP2-depleted cells. The data from the intensity profiles in FIJI were copied to PRISM for graphing.

To quantify midzone microtubule density, a 20-pixel wide line was drawn along the full length of the midzone 30 minutes before complete daughter cell separation, in both control and siCAMSAP2. The mean grey values recorded along the line were then averaged to produce a proxy measure of microtubule density for each condition. For the intensity plot profiles, a line was drawn at the abscission cut event for controls and after microtubule disassembly for siCAMSAP2. The mean grey values along the line were background subtracted for both conditions.

To determine the duration of mitotic events in live long-term image sequences, the DNA signal was used to obtain the timing for chromosome alignment at the metaphase plate and anaphase. Nuclear envelope breakdown (NEBD) was defined as the frame before the elimination of barriers around the nucleus. Metaphase was defined as the frame when chromosomes were aligned at the metaphase plate. Anaphase was defined as the first frame when chromosome separation was detected. We measured NEBD to metaphase and anaphase for both control and siRNA treated cells. To determine the duration of NEBD to metaphase and anaphase, the individual timestamps were recorded and later subtracted from the NEBD timestamp for both control and siRNA treated cells.

To determine the timing of abscission, microtubule signal was monitored as cells progressed through mitosis until a visible microtubule cut was observed from either side of the cell. Z-stacks were used to confirm that the cut represented true severing rather than an out-of-focus artifact. In siRNA treated cells, no clear was observed; instead, the timepoint at which the microtubule visibly dissolved was used as a proxy for separation time.

### Statistical analysis

Measurements were performed in FIJI and statistical analysis was performed in Prism (GraphPad Prism version 9.4.1 for MacOS, (GraphPad Software, San Diego, California USA, www.graphpad.com). The graphs of the data were compiled using Prism (GraphPad Prism version 9.4.1 for MacOS, (GraphPad Software, San Diego, California USA, www.graphpad.com). Images were assembled in Adobe Illustrator (Adobe Inc., version 29.4 Adobe Illustrator).

## Acknowledgements.

We thank Dr. Thomas Maresca and Dr. Lillian Fritz-Laylin for insightful comments on this work. Thank you to the Maresca lab and Fritz-Laylin lab for reagents. Dr. Jacob Ritz provided advice on long-term imaging data and analysis. Funding was obtained from NSF MCB 2134215 and C.F.R was supported by Spaulding-Smith STEM Fellowship from the Graduate School at the University of Massachusetts Amherst.

## Abbreviations

Kif2a: kinesin family member 2A
CAMSAP2: calmodulin-regulated spectrin association protein 2
MAPs: microtubule-associated proteins
ESCRT: endosomal sorting complex required for transport
CHMP2A: charged multivesicular body protein 2A
WT: wild type
AO: anaphase onset
NEBD: nuclear envelope breakdown
PRC1: protein regulator of cytokinesis-1

