## Supplementary figures and images for "Regulation of midzone microtubule dynamics and abscission in human cells by CAMSAP2 and Kif2a"

### Supplemental Figure 1

#### SUPPLEMENTAL FIGURE 1

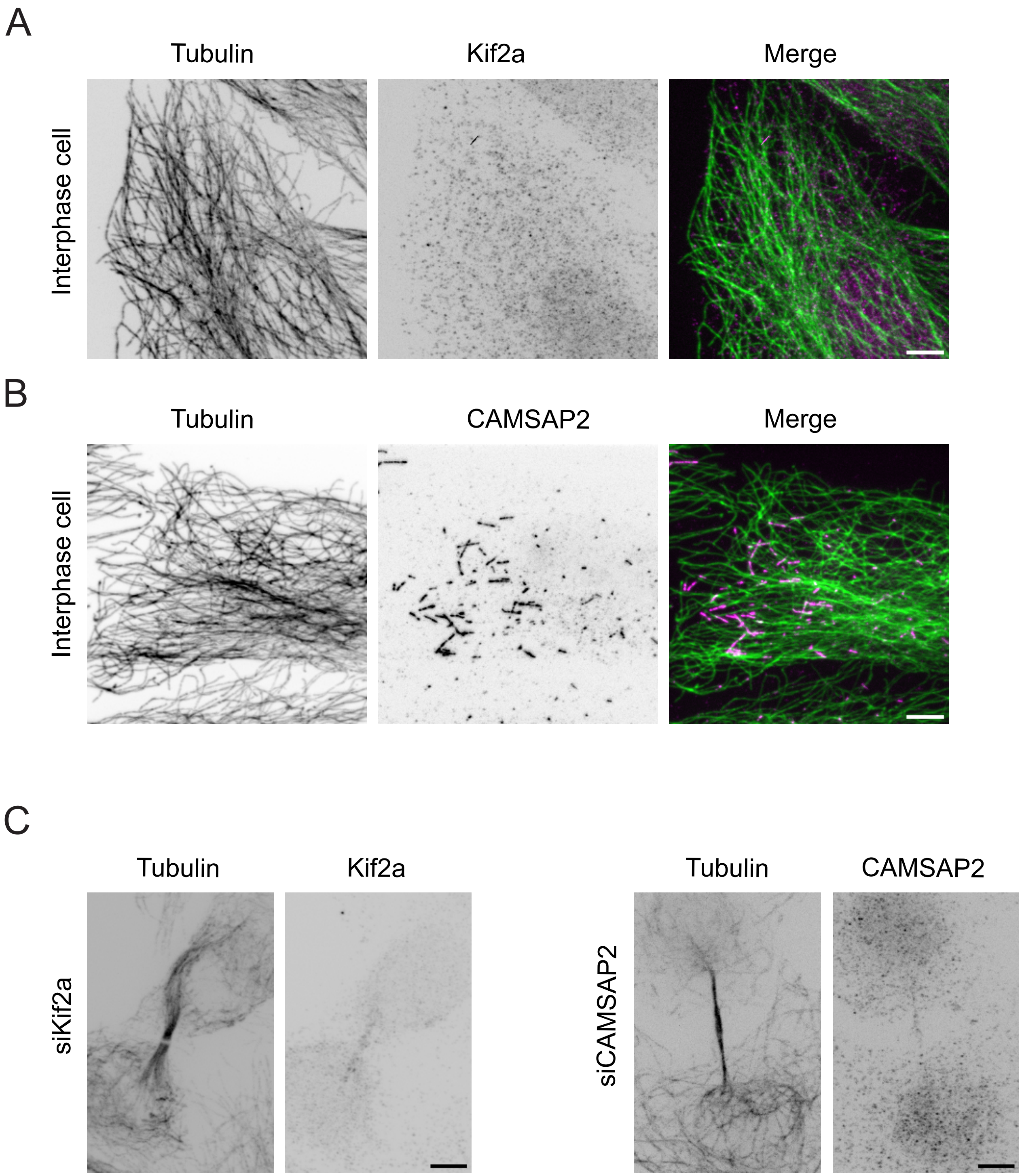

### Supplemental Figure 2

#### SUPPLEMENTAL FIGURE 2

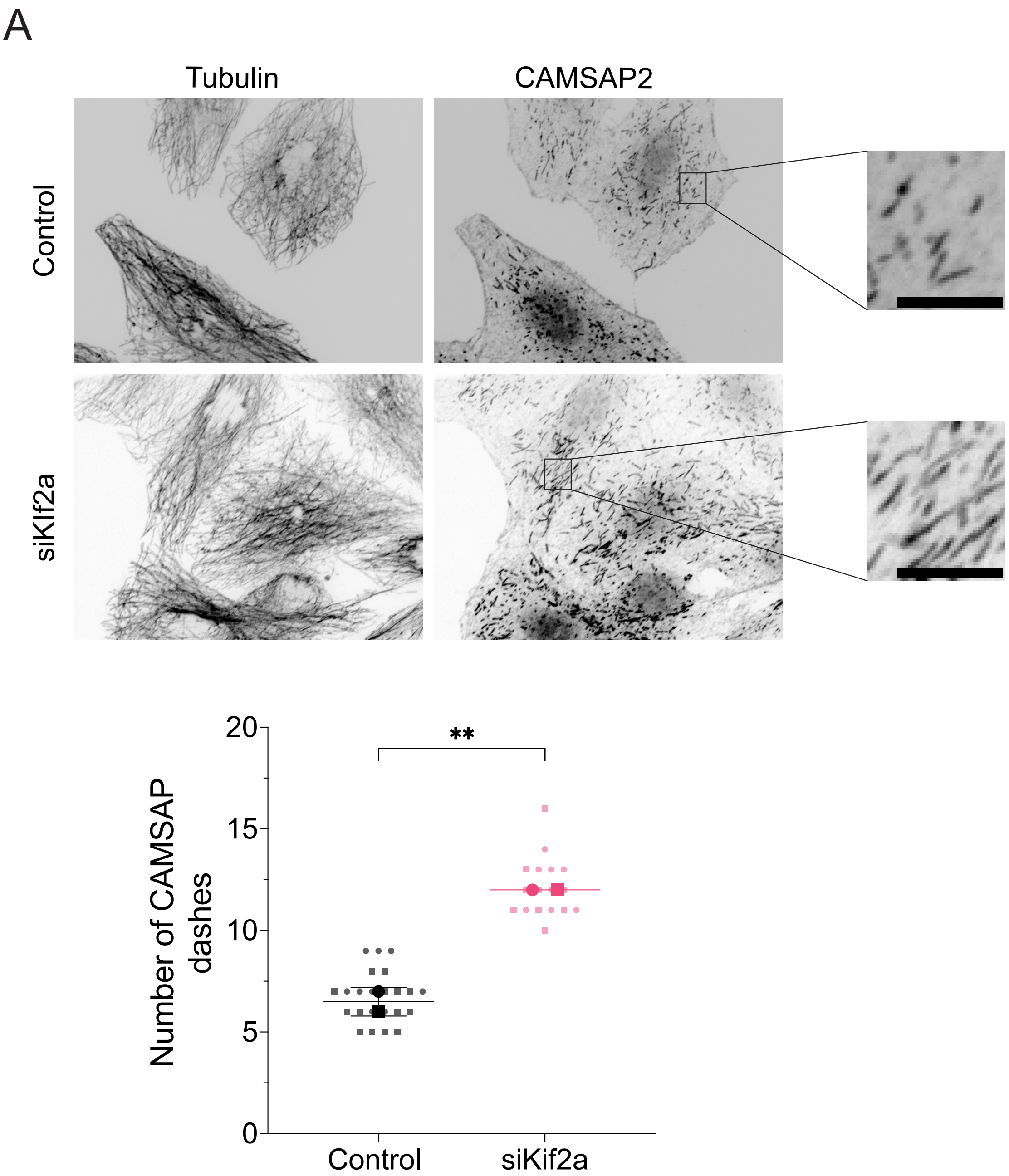

### Supplemental Figure 3

#### SUPPLEMENTAL FIGURE 3

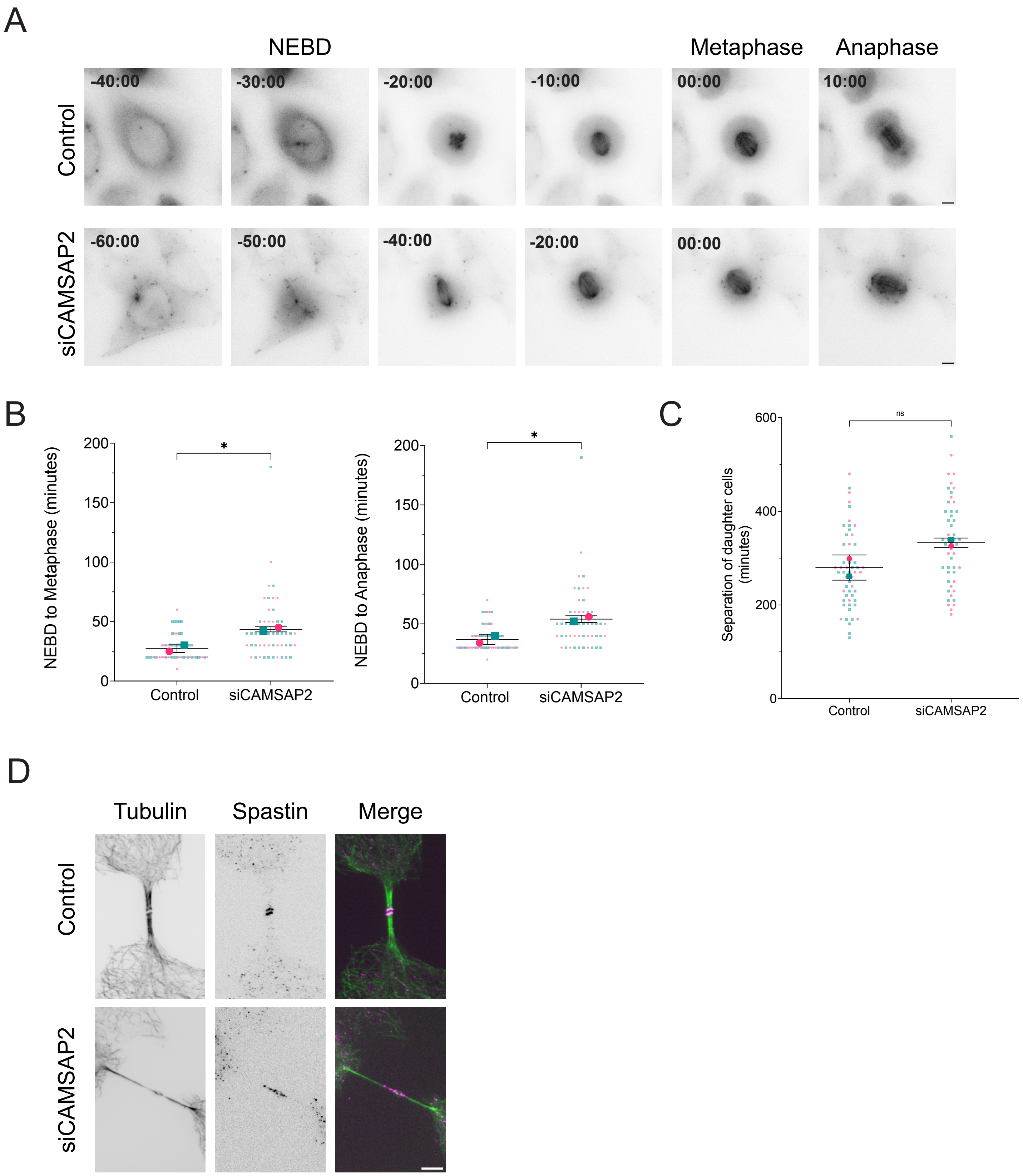
